# Evidence that ion conduction in 5-HT_3A_ receptors proceeds through lateral portals in the cytosol

**DOI:** 10.1101/2020.02.10.943084

**Authors:** Antonia G. Stuebler, Michaela Jansen

## Abstract

The intracellular domain of the serotonin type 3A receptor, a pentameric ligand-gated ion channel, is crucial for regulating conductance. However, the specific ion conduction pathway through this domain is less clear. The intracellular domain starts with a short loop after the third transmembrane segment, followed by a short α-helical segment, a large unstructured loop, and finally the membrane-associated MA-helix that continues into the last transmembrane segment. The MA-helices from all five subunits form the extension of the transmembrane ion channel and shape what has been described as a “closed vestibule”, with their lateral portals obstructed by loops and their cytosolic ends forming a tight hydrophobic constriction. The question remains whether the lateral portals or cytosolic constriction conduct ions upon channel opening. In the present study, we used disulfide bond formation between pairs of engineered cysteines to probe the proximity and mobility of segments of the MA-helices most distal to the membrane bilayer. Our results indicate that the proximity and orientation for cysteine pairs at I409C/R410C, in close proximity to the lateral windows, and L402C/L403C, at the cytosolic ends of the MA-helices, are conducive for disulfide bond formation. While conformational changes associated with gating promote crosslinking for I409C/R410C, which in turn decreases channel currents, crosslinking of L402C/L403C is functionally silent in macroscopic currents. These results support the hypothesis that concerted conformational changes open the lateral portals for ion conduction, rendering ion conduction through the vertical portal unlikely.

**Significance:** The intracellular domain (ICD) of pentameric ligand-gated ion channels (pLGICs) is the most diverse domain within receptors of the Cys-loop superfamily. Despite being the least understood domain of pLGICs, its impact on ion-channel function and contribution to the cytosolic exit pathway of the channel have been investigated. X-ray and cryo-EM structures have captured the structured segments of the ICD of 5-HT_3A_ receptors in different conformational states with lower resolution of the ICD as compared to the other domains. Here, we provide experimentally derived evidence for the importance of the differential mobility of the cytosolic segment of the MA-helices, which supports the existence of lateral portals as opposed to a vertical pathway for 5-HT_3A_ receptors.

## Introduction

5-hydroxytryptamine 3 receptors (5-HT_3_Rs) are pentameric ligand-gated ion channels (pLGICs) in the family of Cys-loop receptors, which also include the cationic nicotinic acetylcholine (nAChR) and the anionic γ-aminobutyric acid (GABA_A_) and glycine receptors. This superfamily is responsible for fast neuronal excitatory and inhibitory neurotransmission in the central nervous system by conducting selected ions across the membrane upon neurotransmitter binding (1,2). The overall structure of pLGICs is conserved. Each subunit consists of three separate domains: the extracellular (ECD), transmembrane (TMD, M1-M4), and intracellular domains (ICD). The ICD of eukaryotic pLGICs, primarily between transmembrane segments M3 and M4, is the most diverse domain with regard to length (50-280 amino acids) and amino acid composition across the family (3).

Successively, structural studies of 5-HT_3A_ (4–7) and nACh (8–11) determined that the ICD consists of the following structural elements: a short post-M3 loop (L1), an amphipathic MX-helix, a long loop (L2) of 60 unresolved amino acids, and the membrane-associated helix (MA-helix), which is continuous with the M4-transmembrane segment (12). The ICD influences the functionality of the receptor: it contributes to desensitization and gating mechanisms, receptor sorting and trafficking by interacting with intracellular proteins, as well as inward rectification and Ca^2+^ permeability (13–20).

Studies have also demonstrated that the ICD can limit the receptor’s single-channel conductance (γ) (21) as it contains the portal(s) for ion access to the cytosol. First structures of the ICD of *Torpedo marmorata* depicted five narrow openings in between adjacent MA-helices, triggering predictions for lateral portals at the TMD-ICD interface (12,22). Structure-function studies based on homology models have since investigated this stretch within the MA-helices and identified specific residues that can influence the receptor’s single-channel conductance (21,23–26). Specifically, three conserved arginines in the 5-HT_3A_ receptor increase the conductance ~33-fold when mutated to their human 5-HT_3B_ subunit counterparts (R432Q/R436D/R440A) (21). Using mutagenesis and Cys scanning/modifications, it was concluded that ions conduct through lateral portals for the triple QDA mutant and that the 5-HT_3A_ receptor has a low single-channel conductance due to steric hindrance and static repulsion by its positively charged residues in the MA-helices (23–25). In reciprocal experiments with α4β2 nAChR, the introduction of arginines at aligned positions reduced the conductance (25). However, the conductance was not fully limited to the characteristically low single-channel conductance of the 5-HT_3A_ homomer, raising the question if other factors are involved in 5-HT_3A_’s 0.6 pS conductance.

More recently, with additional high-resolution structural data becoming available, two alternate exit pathways for ions through the 5-HT_3A_ receptor became apparent: one through the five lateral portals, and another through a vertical portal in continuation with the pore of the transmembrane channel (Fig S1C) (4,6). Although the vertical portal is equipped with a much larger pore diameter (4.2 Å) in the crystal structure, the inside of the pore is lined with hydrophobic residues, plugging the cytosolic end (6,8). The lateral portals, on the other hand, are obstructed by positively charged residues and the L1-loop threading through them (5,6). Clearly, in order to elucidate the ion pathway through the channel, conformational changes of the ICD have to be investigated further.

**Figure 1.**
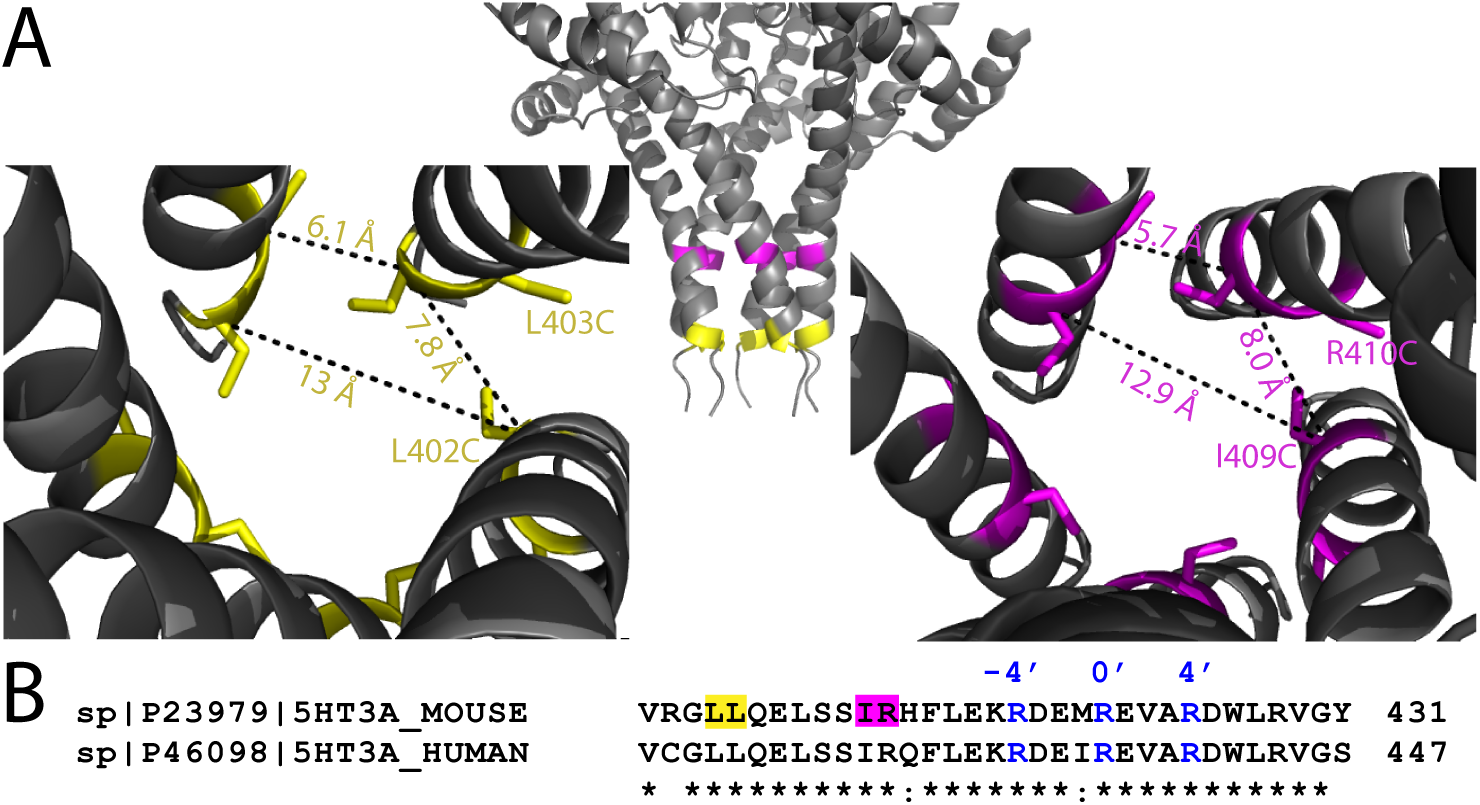
Distances between cysteine pairs engineered in MA-helices in 5-HT_3A_Rs measured between adjacent and non-adjacent subunits. (**A**) (middle panel) Cartoon representation (PDB: 4PIR) of 5-HT_3A_-ICD viewed parallel to the membrane. Yellow (L402/L403) and magenta (I409/R410) residues denote positions where Cys pairs were introduced. (left and right panel) Cartoon representations of the ICD of 5-HT_3A_Rs, viewed down the pore, showing L402C/L403C (yellow, left) and I409C/R410C (magenta, right) residues and the distances between their α-carbons. **(B)** Sequence alignment of the MA-helices of mouse and human 5-HT_3A_Rs with the three arginines mutated to QDA indicated in blue (R-4’, R0’, R4) and the residues mutated to Cys highlighted in yellow (L402/L403) and magenta (I409/R410), respectively.

The overall movement of the receptor entails a global twisting of the ECD and TMD around the central pore (counterclockwise and clockwise for ECD and TMD, respectively), similar to other pLGICs (27,28). Much of this information was first described for the prokaryotic homologues from *Gloeobacter violaceus* and *Erwinia chrysanthemi* (29–31). These receptors could be captured in the open and closed states but lack the ICD only found in eukaryotic pLGICs. The exact conformational changes of the ICD remain largely unexplored but it is known that the ICD undergoes agonist-mediated structural changes, and that the MA stretch may be a mobile structure (32). The flexibility of the MA-helices was even shown to be a determining factor in controlling single-channel conductance in 5-HT_3A_ receptors (33).

A recent cryo-EM structure captured the 5-HT_3A_-ICD in its presumably open state with the lateral portals in a conformation conducive for ion translocation (4). In this structure, the TMD-ICD interface cavity is enlarged by a helical break within the upper section of the MA-helix and the MX-helix is moved outward and upward towards the membrane (Fig S1B).

Previously, we have shown that the ICD alone assembles into pentamers (34) and that disruption of salt-bridges in conjunction with the QDA substitution also abolishes assembly of the ICD into pentamers (35). In the present study, we investigated the mobility of the lower end of the MA-helices (L402-R410, Fig 1), a segment of the MA stretch that remains largely unexplored. This segment starts two α-helical turns below the lowest of the three arginines, −4’ or R416 (mouse 5-HT_3A_, equivalent to R432 in the human sequence) and extends for about two more helical turns (Fig.1). We engineered cysteine pairs three and five helical turns below the lateral portals, and used disulfide trapping to probe the movement at two different levels of the ICD. Through our experiments, we demonstrate that mobility of the MA-helices just below the lateral portals is required to open the cytosolic ion access pathway. On the contrary, inhibiting mobility at the very bottom of the MA-helices, in closer proximity to the vertical opening, had no functional effect on ion-conduction at the macroscopic level.

## Materials and Methods

### Materials

Copper(II) sulfate pentahydrate (Sigma-Aldrich, St. Louis, MO); 1,10-Phenanthroline (Sigma-Aldrich, St. Louis, MO); NEM (*N*-Ethylmaleimide, Sigma-Aldrich, St. Louis, MO); DTT (dithiothreitol, Fisher Scientific, Fair Lawn, NJ); EGTA (Ethyleneglycol-bis(β-aminoethyl)-N,N,N’,N’-tetraacetic acid tetrasodium ≥97%, Sigma-Aldrich, St. Louis, MO); 5-HT (serotonin hydrochloride, Sigma-Aldrich, St. Louis, MO). Stock of serotonin (2 mM), copper(II) sulfate (100 mM), and dithiothreitol (DTT, 1 M) were prepared in distilled water. 1,10-Phenanthroline (0.5 M) was dissolved in dimethyl sulfoxide (DMSO). All solutions, including the Cu:Phe solution, were made in oocyte ringer buffer (OR-2) immediately before experiments were conducted.

### Molecular Biology

Cysteines were engineered into the mouse 5-HT_3A_ receptor (AAT37716) in the pGEMHE vector for oocyte expression (36) that contains a V5-tag (GKPIPNPLLGLDSTQ) close to the N-terminus (16). The desired base pairs coding for Cys were introduced using overlapping primers with the QuikChange II site-directed mutagenesis kit (Agilent Technologies) and were confirmed by DNA sequencing (GENEWIZ, South Plainfield, NJ). The amino acid sequence numbering used corresponds to the numbering of the mature sequence as published with the X-ray structure (6). There are three endogenous cysteines in the ICD of 5-HT_3A_, two in the L2-loop and one ~40 Å apart from the engineered cysteine residues in the X-ray structure (6). cDNA was linearized with the *Nhe*I restriction enzyme and *in vitro* transcribed with the T7 RNA polymerase kit (mMESSAGE mMACHINE^®^ T7 Kit; Applied Biosystems/Ambion, Austin, TX). cRNA was purified with the MEGAclear™ Kit (Applied Biosystems/Ambion, Austin, TX) and precipitated using 5 M ammonium acetate. The cRNA was subsequently dissolved in nuclease-free water and stored at −80 °C.

### *Xenopus laevis* oocyte preparation and injection

*Xenopus laevis* (*X. laevis*) oocytes were isolated, enzymatically defolliculated, and stored as previously described (37). *X. laevis* were handled and maintained following procedures approved by the local animal welfare committee (Institutional Animal Care and Use Committee, IACUC #: 08014, PHS Assurance # A 3056-01). Before injection, oocytes were washed in oocyte ringer buffer (OR-2: 115 mM NaCl, 2.5 mM KCl, 1.8 mM MgCl_2_, 10 mM HEPES, pH 7.5) and maintained in standard oocyte saline medium (SOS: 100 mM NaCl, 2 mM KCl, 1 mM MgCl_2_, 1.8 mM CaCl_2_, 5 mM HEPES, pH 7.5, supplemented with 1% Antibiotic-Antimycotic (100X, 10,000 units/mL of penicillin, 10,000 mg/mL of streptomycin and 25 mg/mL amphotericin B, Gibco, Thermo Fisher Scientific, 5% horse serum) for up to 7 days at 16 °C. Oocytes were microinjected with 10 ng of *in vitro* synthesized cRNA (200 ng/μL) using an automatic oocyte injector (Nanoject II™; Drummond Scientific Co., Broomall, PA). Depending on the construct injected, sufficient expression could be seen 24-48 hrs after injection.

### Electrophysiology

Two-electrode voltage clamp recordings were conducted 1-3 days after injection. The currents were recorded and amplified using a TEV-200A amplifier (Dagan Instruments, Minneapolis, MN), a Digidata 1440A data interface (Molecular Devices, Sunnyvale, CA), and a MiniDigi 1B (Molecular Devices), all controlled by pClamp 10.7 software (Molecular Devices). The oocytes were perfused with OR-2 using gravity flow at an approximate rate of 5 mL/min in a 250 μL chamber. The reagents (Cu:Phe, DTT, and EGTA) were dissolved in OR-2 and perfused in the same manner. All experiments were performed at room temperature (RT, 22–24°C) and a holding potential of −60 mV. The glass pipettes were filled with 3 M KCl and had a resistance of below 2 MΩ. Currents were evoked by 3 μM 5-HT, unless otherwise stated, and the agonist was applied until a stable response was observed to record a maximal current response. The EC_50_ for QDA-I409C/R410C was determined in the presence of 10 mM DTT to minimize disulfide bond formation.

### Oocyte Protein Isolation

*X. laevis* oocytes expressing 5-HT_3A_ and Cys constructs were incubated in 100:200 μM Cu:Phe for 2 min before being homogenized. Uninjected oocytes were treated similarly as control. A crude membrane protein fraction was isolated as described before (38). In brief, oocytes were homogenized using a 200 μL pipette tip in ice-cold vesicle dialysis buffer (VDB: 100 μL/oocyte: 96 mM NaCl, 0.1 mM EDTA, 10 mM MOPS, 0.02% NaN3, 2 mM NEM, pH 7.5), supplemented with protease inhibitor cocktail III (PI, 2 μL/mL; Research Products International, Mt. Prospect, IL), and centrifuged (800 g for 10 min at 4 °C). The supernatant was collected and proteins were obtained by high-speed centrifugation (39,000 g for 1 h at 4 °C). The crude membrane pellet was resuspended in VDB without NEM (3 μL/ oocyte), supplemented with PI cocktail, and stored at −80 °C.

### Western Blotting

For SDS-electrophoresis, proteins isolated as crude membranes as described above, were separated using 4–15% precast gradient TGX Stain-Free™ gels (Bio-Rad) for 35 min at 200 V. Proteins were transferred to polyvinylidene difluoride (PVDF) membranes using the Trans-Blot Turbo Transfer System (Bio-Rad). Membranes were blocked with 5% blotting-grade blocker (Bio-Rad) in Tween-tris-buffered saline (TTBS, 100 mM Tris, pH 7.5, 0.9% NaCl, 1% Tween-20) [25 – 30 mL/membrane]) overnight. The membranes were incubated with the primary V5 HRP-conjugated antibody (1:5,000, V5-HRP antibody, Invitrogen) in TTBS with 5% blotting-grade blocker overnight at 4 °C. After the removal of the primary antibody, the membranes were washed 5 times for 5 min with TTBS. Proteins were visualized by chemiluminescent detection (ImageQuant LAS 4000, GE Healthcare Life Sciences) of peroxidase substrate activity (SuperSignal West Femto Maximum Sensitivity Substrate, Thermo Scientific). Band intensities were quantified using UN-SCAN-IT gel™ (Silk Scientific, Inc.: Orem, UT). The DTT-resistant band (running at ~130 kDa) was excluded from the quantification but caused the background of the membrane to appear darker in close proximity to the location of the dimer, potentially increasing the band intensity.

### Data Analysis

Data was analyzed using pClamp, Origin (OriginLab Corporation: Northampton, MA), Prism 6 Software (GraphPad Prism^®^; La Jolla, CA), and UN-SCAN-IT *gel*™ (Silk Scientific, Inc.: Orem, UT). Data is shown as mean ± S.D. (n≥3) with the maximal current induced by 5-HT as the normalizing standard (100% of current response) for other current amplitudes recorded in the same oocytes. Statistical significance was determined with unpaired t-test, paired t-test between the initial inward current and each set of conditions or one-way ANOVA, Dunnett’s multiple comparisons test (*p≤0.01 unless otherwise stated). All figures and graphs were made in Origin, PyMOL Molecular Graphics System (Schrödinger, LLC), and Adobe Illustrator CC 2018.

## Results

### Design of engineered cysteine pairs, L402C/L403C and I409C/R410C, to crosslink MA-helices in 5-HT_3A_Rs

To investigate the mobility of the cytosolic end of the intracellular domain of 5-HT_3A_R, we engineered two different cysteine pairs below the proposed lateral portals at different levels in the ICD, almost two helical turns apart: L402C/L403C (Fig. 1A, yellow) and I409C/R410C (Fig.1A, magenta). For each double-Cys construct, the mutated cysteines are adjacent to one another on the same subunit (Fig. 1), with side chains extending towards the cysteines on adjacent subunits. Given that the average Cα-Cα’ carbon distance for the engineered Cys in adjacent subunits is similar to the average distance found between disulfide-bonded cysteines of 5.6 Å (39,40), cysteines placed at these locations should enable crosslinking of MA-helices (Table 1). Both the orientation of the side chains and α carbon separation predispose these Cys pairs for disulfide bond formation. All other side chain orientations or distances in between these positions are less suited for disulfide bond formation, with the next closest distance of 6.4 Å between E404 and L406. Double-cysteine pairs were introduced in the 5-HT_3A_- and QDA-background to compare the impact of MA-flexibility in low- (5-HT_3A_) and high- (5-HT_3A_-QDA) conductance receptors.

**Table 1:** Distances (in Å) between engineered cysteines and pore diameters in different X-ray and cryo-EM structures, measured in Pymol.

| PDB | L402C/L403C |  |  | L409C/L409C |  |  |
| --- | --- | --- | --- | --- | --- | --- |
|  | Across the pore | L402C-L403C | L402C-L402C | Across the pore | I409C-I410C | I409C-I409C |
| 4PIR x-ray | 13.0 | 6.1 | 7.8 | 12.9 | 5.7 | 8.0 |
| 6BE1 (Apo-closed) | 14.7 | 7.3 | 9.4 | 13.6 | 7.4 | 9.1 |
| 6HIS (T inhibited) | 14.6 | 6.9 | 9.0 | 14.2 | 6.3 | 8.8 |
| 6HIQ (I2 intermediate) | 14.8 | 6.9 | 8.9 | 13.6 | 6.0 | 8.6 |
| 6DG8 (State 2 open) | 15.4 | 7.7 | 9.0 | 15.3 | 8.3 | 9.5 |

### L402C/L403C and I409C/R410C Cys pairs form intersubunit disulfide bonds as visualized in western blots after SDS-PAGE separation

To evaluate whether each Cys pair promotes intersubunit crosslinking (i.e. disulfide bridge formation between MA-helices), we compared the oligomeric state/s induced by oxidizing and reducing conditions, as well as by application of the agonist, of each construct to the background receptor (5-HT_3A_ or 5-HT_3A_-QDA) by western blot after SDS-polyacrylamide gel electrophoresis (Fig. 2). Constructs were expressed in *X. laevis* oocytes that were exposed to agonist (3 μM 5-HT) or oxidizing agent for 2 min before they were homogenized to generate crude membrane protein fractions. To promote an oxidizing environment and encourage disulfide bond formation in the Cys-pairs, we exposed oocytes expressing 5-HT_3A_ constructs to 1,10-phenanthroline in the presence of copper (Cu:Phe, 100:200 μM) for 2 min. Cu:Phe is an oxidizing agent that has been widely used as a catalyst for oxidation of sulfhydryl groups to disulfides (41–44), and its oxidative mechanism involves diffusible reactive intermediates that can penetrate protein interiors (45–47). Each condition used sixteen or more oocytes. 5-HT_3A_ channels were detected via a V5-epitope tag located near the N-terminus (Fig. 2A). The band intensities varied with different expression levels and were adjusted accordingly for each lane. All 5-HT_3A_ receptors have monomer bands at ~55 kDa (Fig 2A), which is equivalent to the theoretical molecular weight of the monomer. There is also a second band present in all lanes (band indicated by *), which has previously been observed upon extraction of the channel from oocytes and runs at a slightly higher molecular weight (~130 kDa) than the expected dimer (~110 kDa, indicated by <).

**Figure 2.**
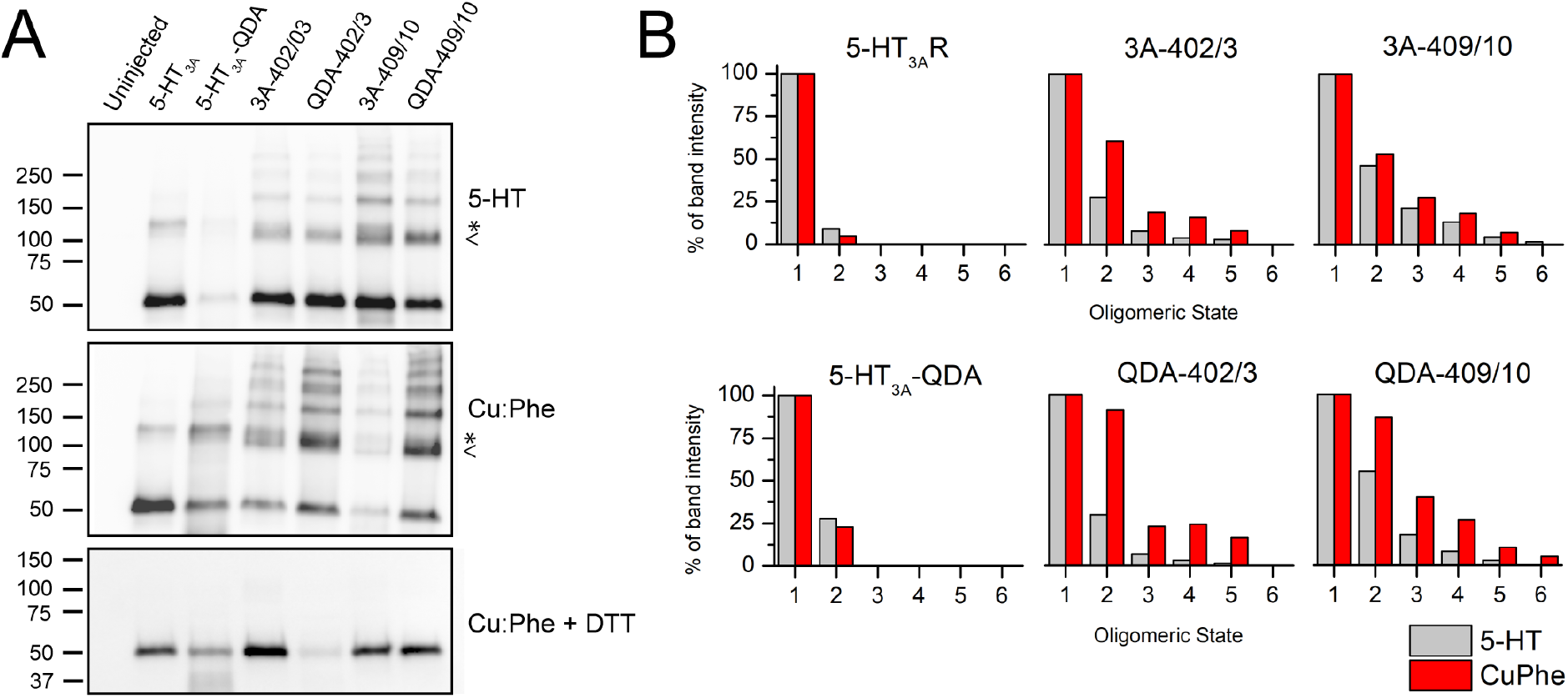
Agonist- or Cu:Phe-application induce disulfide bond formation for L402C/L403C and I409C/R410C expressed in *Xenopus* oocytes. **(A)** 5-HT_3A_ and construct-expressing *X. laevis* oocytes were treated to mimic the electrophysiological experiments (either exposed to 2 min of 3 μM 5-HT or 100:200 μM Cu:Phe). Crude membrane fractions were separated by SDS-PAGE and loading was adjusted for construct-dependent expression levels. 5-HT_3A_ subunits were detected after western blotting. (**A**) Membrane fractions from oocytes treated with the agonist 5-HT (top), and the oxidizing agent Cu:Phe (middle) were run without reducing agent. A monomer band (55 kD) is observed for all constructs, and higher crosslinked oligomeric states for all double-Cys mutants. Note, a band (indicated by *) running at a slightly higher molecular weight than dimers (indicated by <) is seen in all samples (~130 kDa), including in the absence of engineered Cys. Oxidized membrane fractions were also separated in sample buffer with reducing agent (DTT) to show release of the crosslink through disulfide reduction. **(B)** The band intensity was quantified using UN-SCAN-IT *gel*™. Monomer bands were used as control (100%) for each construct with each condition. The background band at a slightly higher molecular weight than dimer present in all 5-HT_3A_-expressing samples leads to some “dimer” signal in the WT and QDA. Overall Cu:Phe (red bars) increases the prevalence of crosslinked species compared to agonist (grey bars), which appears to be more pronounced in the QDA background.

Larger bands (>150 kDa) are visible for each L402C/L403C and I409C/R410C Cys pair, indicating that these receptors form higher oligomeric states in the presence of 5-HT or Cu:Phe (Fig. 2A). The presence of these higher molecular weight bands was completely reversed by reduction of the Cu:Phe crude membrane fractions with DTT, suggesting that they represent oligomers stabilized by disulfide bridges (Fig. 2A bottom panel). Additionally, the different band intensities were quantified using UN-SCAN-IT *gel*™ and are shown as percentages normalized to the monomer band set to 100% for each oligomeric state and each construct (Fig. 2B). While the dimer band is prominently present in all Cys pairs, higher oligomeric bands have a larger intensity in the QDA-background of the Cys pairs as compared to the regular 5-HT_3A_-background (Fig. 2B).

### Agonist-induced crosslinking observed in macroscopic currents

To test functionality of double-Cys receptors and the stability of their current responses, we measured 5-HT induced currents from *X. laevis* oocytes expressing 5-HT_3A_, 5-HT_3A_-QDA, and each Cys-pair using two-electrode voltage clamp recordings (Fig. 3A). All receptors exhibited inward currents in response to the application of 3 μM 5-HT, a concentration ~5-fold above EC_50_ values for 5-HT_3A_, QDA, and QDA-I409C/R410C (16), indicating that double-Cys channels form functional receptors. Brief and repeated 5-HT exposures demonstrated that the inward current amplitudes of 5-HT_3A_, 5-HT_3A_-QDA, and double-Cys pair L402C/L403C were stable (variations <10%, Fig 3A).

**Figure 3.**
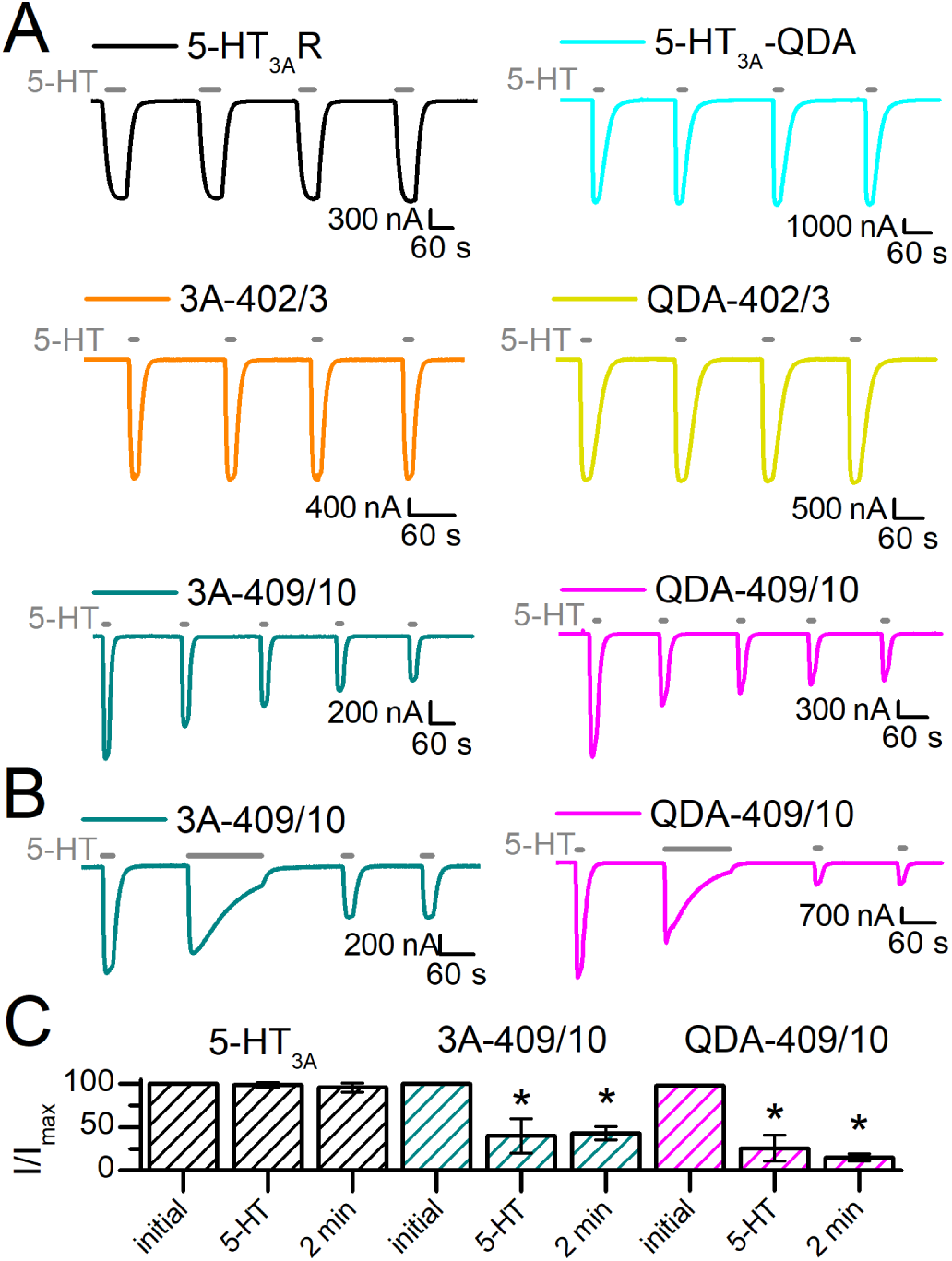
Agonist-induced crosslink of the double Cys pair I409C/R410C induced by agonist application(s). **(A)** Sample traces of currents recorded from *X. laevis* oocytes expressing 5-HT_3A_ wildtype and engineered constructs in response to 3 μM 5-HT (grey bars). 5-HT_3A_R (black), 5-HT_3A_-QDA (teal), 5-HT_3A_-L402C/L403C (orange), and QDA-L402C/L403C (yellow) show stable responses to repeated applications of the agonist, with both the current amplitudes and kinetics unaltered. 5-HT_3A_-I409C/R410C (green) and QDA-I409C/R410C (magenta) show successively decreasing current amplitudes with each 5-HT-application until a stable response is reached. **(B)** Sample traces of 5-HT_3A_-I409C/R410C (green) and QDA-I409C/R410C (magenta) in response to 3 μM 5-HT (grey bars). The initial inward current represents the reference current amplitude (100% of current) and is followed by an agonist application of 2 min. After this prolonged 5-HT exposure, both Cys pairs exhibit reduced but stable inward currents. **(C)** Quantification of the reduced current amplitudes after repeated (5-HT) and prolonged (2 min) 5-HT application(s) is also shown in bar graphs. Data is shown as mean±S.D. (n≥3). Statistical significance was determined with paired t-test between the initial inward current and the stable, reduced current amplitude after agonist exposure(s) (*p≤0.01).

In contrast to the other receptors, the Cys-pair I409C/R410C in both backgrounds showed current attenuation with repeated 5-HT application (Fig. 3A, bottom traces) until a stable level was reached, ~38% and ~27% of the original current, for 5-HT_3A_-I409C/R410C and QDA-I409C/R410C, respectively. This decrease in current was not only induced by repeated agonist applications. I409C/R410C receptors also showed profound current attenuation of test currents following a single longer 5-HT exposure of 2 min (Fig. 3B). The current reduction after repeated agonist applications was not significantly different than the current attenuation with a longer single 5-HT exposure (unpaired t-test, p>0.05). The same experimental setup yielded desensitization of the majority of 5-HT_3A_ channels during the 2-min agonist application, but no reduction in test current amplitudes of subsequent 5-HT applications (Fig. 3C, black bars).

For channels with only a single engineered Cys, 5-HT_3A_-I409C and QDA-I409C, inward current attenuation was not observed (Fig. S2). On the contrary, these channels showed stable responses to agonist application, comparable to 5-HT_3A_ and QDA without engineered Cys.

### Effects of Cu:Phe, DTT, and EGTA on 5-HT-induced currents

Oocytes expressing 5-HT_3A_ constructs were treated with the oxidizing agent Cu:Phe (100:200 μM) for 2 min (Fig. 4A, Fig. S3A), following the initial inward current response evoked by 5-HT (42,43,48). This first inward current was tested for stability in 5-HT_3A_Rs and all constructs, except in 5-HT_3A_-I409C/R410C and QDA-I409C/R410C, due to the current decrease observed with repeated agonist application (Fig. 3A). Following the 2 min exposure to Cu:Phe, the oocytes were washed for 6 min with OR-2 before another agonist-induced current was recorded (Fig 4A, second inward current) and tested for stability. Lastly, we investigated the effect of applying 10 mM DTT (Fig. 4A, Fig. S3A), a reducing agent capable of converting protein disulfides to sulfhydryls via thiol-disulfide exchange reaction, or 1 mM EGTA (Fig. 4B, Fig. S3B), a copper chelating agent (49). DTT or EGTA was applied to the same oocyte after Cu:Phe exposure for 2 min, followed by a 6 min OR-2 wash before another 5-HT application (final inward current, DTT: Fig 4A, EGTA: Fig 4B).

**Figure 4.**
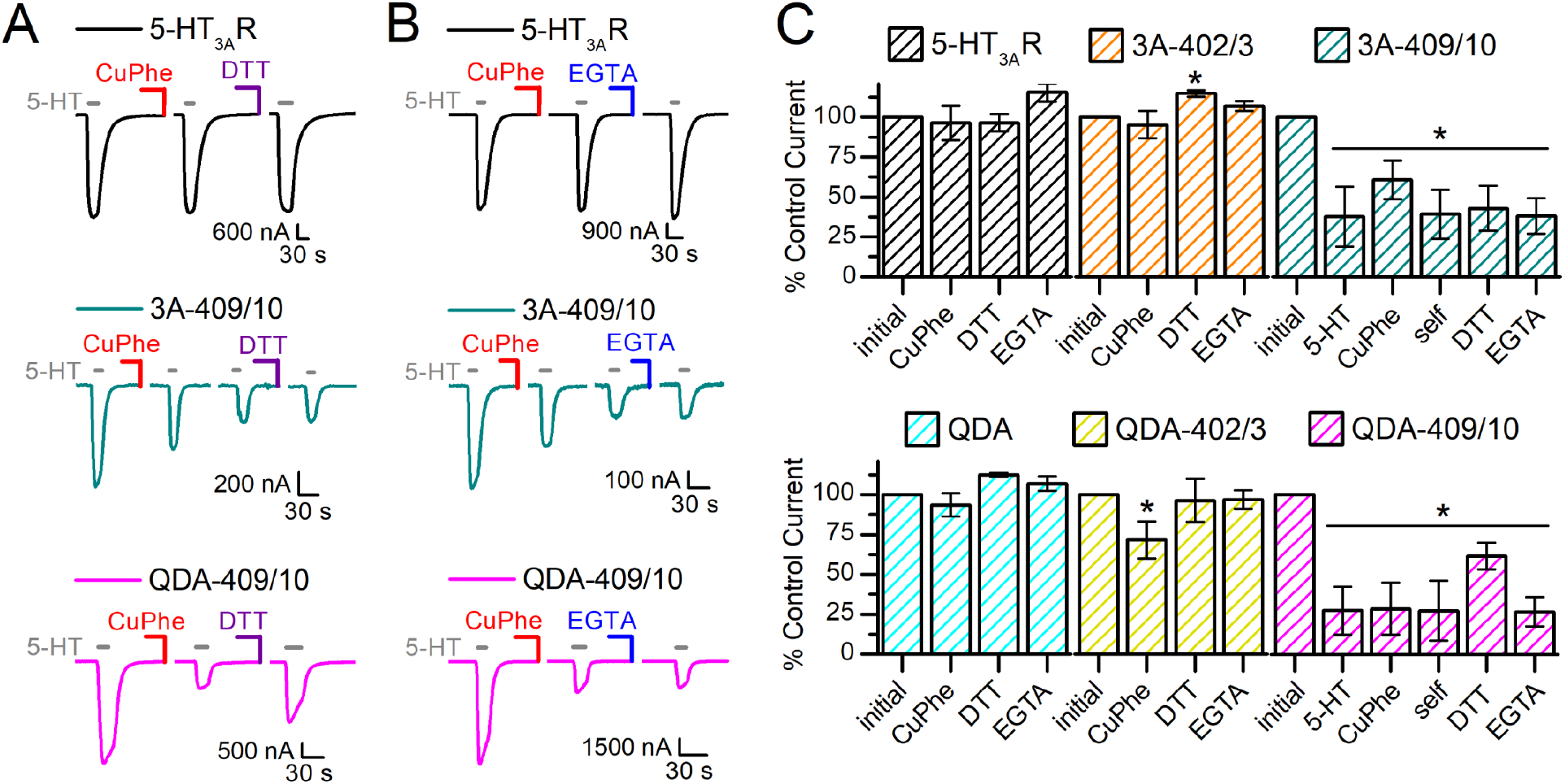
Effect of oxidizing (Cu:Phe), reducing (DTT), and chelating (EGTA) agents on 5-HT_3A_ and I409C/R410C Cys pairs. **(A)** Sample traces of 5-HT_3A_R (black), 3A-I409C/R410C (green), and QDA-I409C/R410C (magenta). The inward currents were evoked with 3 μM 5-HT. The initial inward current represents the reference current amplitude (100% of current). Following the 5-HT application(s), the oocytes were exposed to 2 min of Cu:Phe (100:200 μM) and then a 6 min wash with OR-2 (not pictured) before another application of 5-HT (second inward current depicted). 5-HT was applied until a stable current response was achieved (stable response to 5-HT is the inward current before DTT, labeled “5-HT” in **(C)** for I409C/R410C mutants). Lastly, 10 mM DTT was applied to the oocytes for 2 min, followed by a 6 min OR-2 wash (not pictured) and 5-HT (last inward current pictured). **(B)** Sample traces of 5-HT_3A_R (black), 5-HT_3A_-I409C/R410C (green), and QDA-I409C/R410C (magenta). Same experimental set-up as in **(A)** but during the last step 1 mM EGTA was applied instead of DTT. **(C)** Quantitative representation of current amplitudes, including Fig. S3. The initial inward current represents 100% of the current response. Data is shown as mean±S.D. (n≥3). Statistical significance was determined with one-way ANOVA, Dunnett’s multiple comparisons test between the initial inward current and each set of conditions (*p≤0.01).

Overall, 5-HT-induced inward currents of 5-HT_3A_ and 5-HT_3A_-QDA were unaffected by Cu:Phe, DTT, and EGTA (Fig. 4). The current amplitudes of 5-HT_3A_-L402C/L403C showed no change after Cu:Phe or EGTA, however increased by ~14% after exposure to DTT (Fig. S3, Fig. 4C). The same Cys pair in the QDA-background, QDA-L402C/L403C, showed reduced current amplitudes after Cu:Phe (71.7±11.6% of initial, n=12) as compared to the initial current, which could be recovered by either DTT or EGTA (Fig. S3, Fig. 4C). Overall, Cu:Phe had the most pronounced effect on 5-HT_3A_-I409C/R410C and QDA-I409C/R410C, decreasing their inward current amplitudes to ~61% and ~29% of the initial inward current, respectively (Fig. 4). An effect not observed in the single Cys construct, 5-HT_3A_-I409C (Fig. S2). In the case of 5-HT_3A_-I409C/R410C, the current responses after Cu:Phe were not stable, and the amplitudes were further attenuated to ~39% of the original current with repeated agonist applications (Fig 4. middle panel, third inward current evoked by 5-HT in sample traces). This reduction in current response could not be recovered for 5-HT_3A_-I409C/R410C with either DTT or EGTA application and could be partially recovered with DTT (61.6±8.32% of initial current) but not EGTA for QdA-I409C/R410C.

## Discussion

In the present study, we investigated the effect of constraining the cytosolic end of the MA-helices of 5-HT_3A_ by using disulfide bond formation between engineered cysteines. Disulfide trapping is a powerful tool to analyze backbone and domain motions in proteins of unknown and known high-resolution structure with the advantage that these experiments can be carried out in the native environment of the protein (39,46), including within the membrane. We engineered double-Cys pairs within the MA-helix of 5-HT_3A_ subunits with the goal of using intersubunit disulfide bonds between subunits to clamp the MA-helices in a constrained conformation to gain insight into the importance of the movement of the lower end of the MA-helices during gating. The Cys substitutions were introduced two and four helical turns below the lowest arginine (R416 or −4’, Fig 1B) located in the lateral portals, the part of the MA-helix that so far has been orphaned by functional studies. The same Cys-pairs were introduced into the QDA-background that has a ~33-fold increased conductance (21).

SDS-PAGE separation of crude membrane fractions followed by immunoblotting indicated that the MA-helices are crosslinked for both Cys-pairs following treatment with agonist or oxidizing agent, manifested as bands representing dimers and higher-order oligomers (Fig. 2A) (50). Oligomer bands were only observed for the Cys-pairs and not for 5-HT_3A_, QDA or single-Cys constructs (Fig. S2) and could be completely reduced to monomers in the presence of a reducing agent, ascertaining that these bands were indeed the result of disulfide bonds.

Functionally, the double-Cys pairs at I409C/R410C showed agonist-induced disulfide bond formation, indicated by the decrease in current amplitude with repeated or prolonged applications of 5-HT until a steady state was reached (Fig. 3 and 4). A more oxidizing environment mediated by the catalyst Cu:Phe also led to disulfide-bond formation (Fig. 4), which was not stable or complete for this Cys pair in the 5-HT_3A_-background. Agonist-applications reduced the current further for 5-HT_3A_-I409C/R410C but current reductions could neither be recovered with reducing nor chelating agents. For QDA-I409C/R410C on the other hand, crosslinking was partially reversed with DTT but not the chelating agent EGTA, indicating that the reduction in current is caused by disulfide bond formation and not heavy metal binding.

The other Cys pairs at L402C/L403C only showed a small change in current response following agonist and oxidizing agent application. For QDA-L402C/L403C oxidizing reagent-induced current reduction and reductant- and EGTA-induced reversal were observed (Fig. 4C and Fig. S3, yellow). The 5-HT_3A_-L402C/L403C Cys pair only had a small current increase after DTT (Fig. 4C).

Disulfide bond formation of cysteine side chains depends on the presence of an oxidizing environment as well as the sulfhydryl collision frequency, which is influenced by their separation distance, their orientation in the protein, and the backbones mobility in the region (51). Crosslinking in the absence of an oxidizing catalyst, as observed upon agonist-administration for the Cys pairs at I409C/R410C, indicates a relatively high propensity for disulfide bond formation. The reaction mechanism does not differ fundamentally from crosslinking in more oxidizing environments created by Cu:Phe (51,52). In the case of I409C/R410C, agonist application(s) achieved a large and stable current reduction, suggesting that serotonin induces a motion in the receptor that promotes disulfide bond formation between these positions (53). In the presence of 5-HT, the channels undergo transitions between open, desensitized, and closed states. We cannot, with certainty, distinguish in which state agonist-induced disulfide bond formation occurs due to the millisecond timescale of transitioning (42,51).

The disulfide-bond formation initiated by agonist application yields two subsets of information. Firstly, disulfide bonds have well-defined distance and angular constraints, which indicates close proximity for the two Cys when a cystine forms (39). Secondly, disulfide bond formation at I409/R410 puts structural constraints on the mechanism of conformational changes, giving insights into the relationship between structural changes and function (54). There are several factors that could mediate the reduction in current upon crosslinking, including a decrease in single-channel conductance, especially for the QDA-background, a change in open probability, locking the channels in a non-conducting conformation, and altered kinetics. Based on the present data we cannot distinguish between these possibilities.

SDS-PAGE separation and western blot following treatments that favor crosslinking revealed that most of the crosslinked Cys-pairs run as dimers as opposed to higher oligomers (Fig. 2B). We infer that when two adjacent subunits are crosslinked by the formation of one disulfide bond within the pentamer, this disulfide bond constrains the concerted movement also of the other subunits, affecting orientation and backbone mobility of the MA-helices.

Agonist-induced current reduction was absent for the Cys pairs at L402C/L403C (Fig. 3A). A minimal current reduction after Cu:Phe application was observed, that was recovered after either DTT or EGTA incubation, pointing towards a small inhibitory effect by heavy-metal binding (Fig. 4C) as both DTT and EGTA can act as metal chelators. Our results indicate that the crosslinking observed for L402C/L403C using SDS-PAGE separation and immunoblotting is largely functionally silent in macroscopic currents, which could indicate their proximity to each other in the equilibrium structure (46) and that conformational flexibility at the vertical end of the pore is not required for ion conduction.

A characteristic specific to the A subunit of 5-HT3 receptors is that the MA-helices are lined with positively and negatively charged residues that interact to form inter- and intrasubunit salt bridges (33,35). Another notable salt bridge is located between the MA-helix and the L1-loop: D312 at the center of L1 is straddled by the 0’ and 4’ arginine residues (mouse R420 and R424, human R436 and R440) from the MA-helix of the adjacent subunit. Amino acid substitutions introduced in the proximity of the five lateral portals, such as the QDA substitution, lead to the breakage of these salt bridges, which enhances flexibility of the MA-bundle, increases single-channel conductance, and disrupts pentamerization of the ICD (21,33,35). Based on these observations, we hypothesize that partial release of intersubunit salt-bridges in the QDA background yields a more open and unclamped/flexible arrangement of the ICD, in particular the MA-helices. This could explain our experimental differences observed between the 5-HT_3A_ and QDA-backgrounds. The QDA-I409C/R410C Cys pair has a ~10% larger current reduction after the restriction of its increased MA-mobility after 5-HT application as compared to the same Cys pair in the 5-HT_3A_ background while the current reduction is significantly different by about 30% directly after Cu:Phe application between the two backgrounds (unpaired t-test, t(19)=5.21, p=4.96×10^-5^). The overall quantification of band intensities for each construct in each condition revealed that the Cys pairs in the QDA-background show higher percentages for dimers and some higher oligomeric states as compared to the WT-background (Fig. 2B).

QDA-I409C/R410C could also be partially recovered with DTT (Fig. 4C). Based on the mechanism of the involved redox chemistry, reduction with DTT involves a disulfide exchange reaction, necessitating access of the DTT molecule in close apposition to the disulfide bond to be reduced. Therefore, the inability to reverse oxidation with DTT may be due to the reduced accessibility of the less mobile and rigid MA-bundles in the WT background (33) and an increased mobility mediated by the QDA substitutions.

Overall, we conclude that disulfide bonds form for both double-cysteine pairs based on oligomers observed after immunoblotting and our functional data. We infer that conformational changes associated with gating are more pronounced for the upper I409C/R410C pair that sits three helical turns below the frame of the lateral windows and on the upper end of the hydrophobic plug. The area below this plug, where the helices converge, seems to be involved in smaller motions while the ICD above twists around the central pore of the receptor.

These findings can be compared to the mechanics of an umbrella, where the MA-helices represent the rods of the umbrella. In the inverted umbrella the ferrule/top notch corresponds to the cytosolic end of the MA-helices. Low on the converging rods/MA-helices we find L402C/L403C whereas I409C/R410C is positioned towards the top of the rods (Fig. 5). When the umbrella is closed (aka. the receptor is in a nonconducting state) the rods are aligned with the shaft (ion pore), in a vertical line, and both the lower (L402C/L403C) and upper portion (I409C/R410C) of the rods are in close proximity to corresponding adjacent rod positions. When the umbrella opens, the upper rod positions (I409C/R410C) move outward to allow for the canopy to open, while the lower positions closer to the notch (L402C/L403C) undergo smaller movements at the end of the shaft.

**Figure 5.**
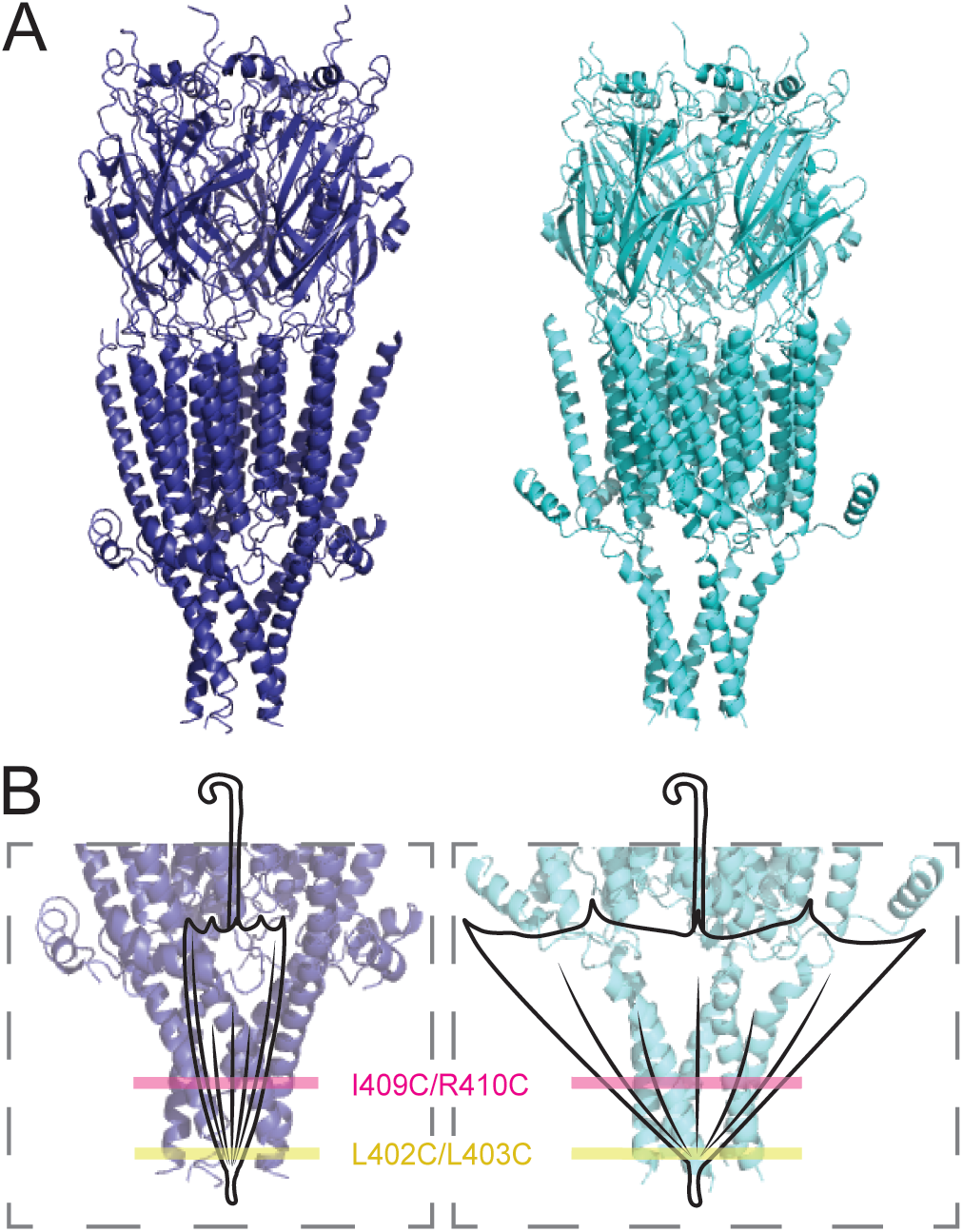
Umbrella analogy of MA-helix mobility. The MA-helices move in a comparable fashion between closed and open states, as do umbrella rods between closed and open states. The L402C/L403C (yellow) Cys pair is located closer to the top notch, where the rods of the umbrella converge, and I409C/R410C (magenta) further away from the ferrule. When the umbrella is closed (the receptor is in a non-conducting state – blue (PDB: 6BE1)), the rods are aligned with the shaft of the umbrella. The rods are stretched away when the canopy opens (receptor in a conducting state – cyan (PDB: 6DG8)). Note that the distance changes between rod positions closer to the ferrule/top notch are smaller and increase with increasing distance to the ferrule.

Structures indicate a clockwise rotational movement of the TMD between closed and open states (4). The TMD is structurally linked to the ICD by the continuation of the M4 transmembrane segment into the MA-helix. This twisting movement of the MA-helices could break the salt bridge network, including D312, which holds the L1-loop in place, resulting in the L1-loop moving outward which in turn allows translation of the MX-helix towards the membrane inner leaflet (Fig. S1).

## Conclusion

Our results investigate the importance of the mobility of the cytosolic end of the MA-helices by using a disulfide chemistry approach. Intersubunit crosslinking through agonist-activation/oxidation three helical turns below the lateral portals leads to smaller macroscopic currents. This indicates that mobility/flexibility of the MA-helices below the lateral portals is required to open the cytosolic ion pathway. In contrast, disulfide bond formation at the cytosolic end of the MA-helices, where the helices converge, is functionally silent in macroscopic currents. The absence of a functional effect of this disulfide crosslinking further supports that ions do not exit the cytosolic vestibule along the long channel axis, but instead, through five lateral portals that open through concerted conformational changes involving MA-helices, L1-loops, and by extension, also MX-helices. Further studies are needed to investigate the concerted mechanism and movements and may lead to the identification of a future therapeutic target as the MA-helices can affect channel function.

## Author Contributions

M.J. and A.G.S. designed research. A.G.S. performed experiments. A.G.S and M.J analyzed data. A.G.S. and M.J. wrote the paper.

## Acknowledgments

We thank the TTUHSC Core Facilities: some of the images and or data were generated in the Image Analysis Core Facility & Molecular Biology Core Facility supported by TTUHSC.

## SUPPORTING INFORMATION

**Figure S1.**
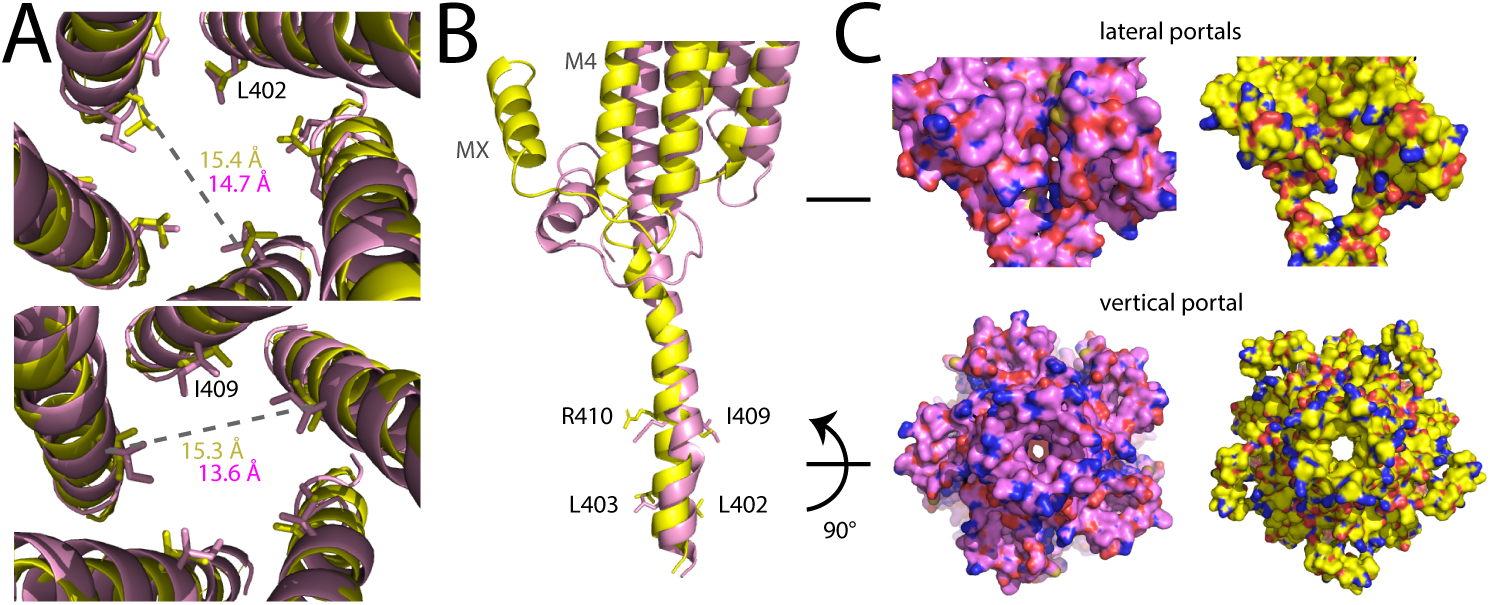
Structural alignment of 5-HT_3A_ Apo (PDB: 6BE1, pink) and State 2 (PDB: 6DG8, yellow) and detailed view of possible ion exit pathways. **(A)** View down the pore of the ICD of 5-HT_3A_. Side chains visible for L402/L403 (top panel) and I409/R410 (bottom panel) as well as the pore diameter for the different structures. **(B)** Alignment of a single subunit of 5-HT_3A_, cartoon representation, emphasizing the differences between the open and closed conducting states. (C) Surface representation of 5-H_3A_ in different conformational states, showing a detailed view of the plausible ion exit pathways. (top panel) Detailed view of the lateral portals between two adjacent MA-helices around the R0’ (R436), viewed parallel to the membrane. State 2, which is potentially conducting (PDB: 6DG8), shows a larger lateral opening as compared to the apo structure (PDB: 6BE1). (bottom panel). Receptors viewed from the intracellular side, comparing the diameter of the vertical pore in the different structures, again with state 2 having a slightly larger aperture as compare to the apo state.

**Figure S2.**
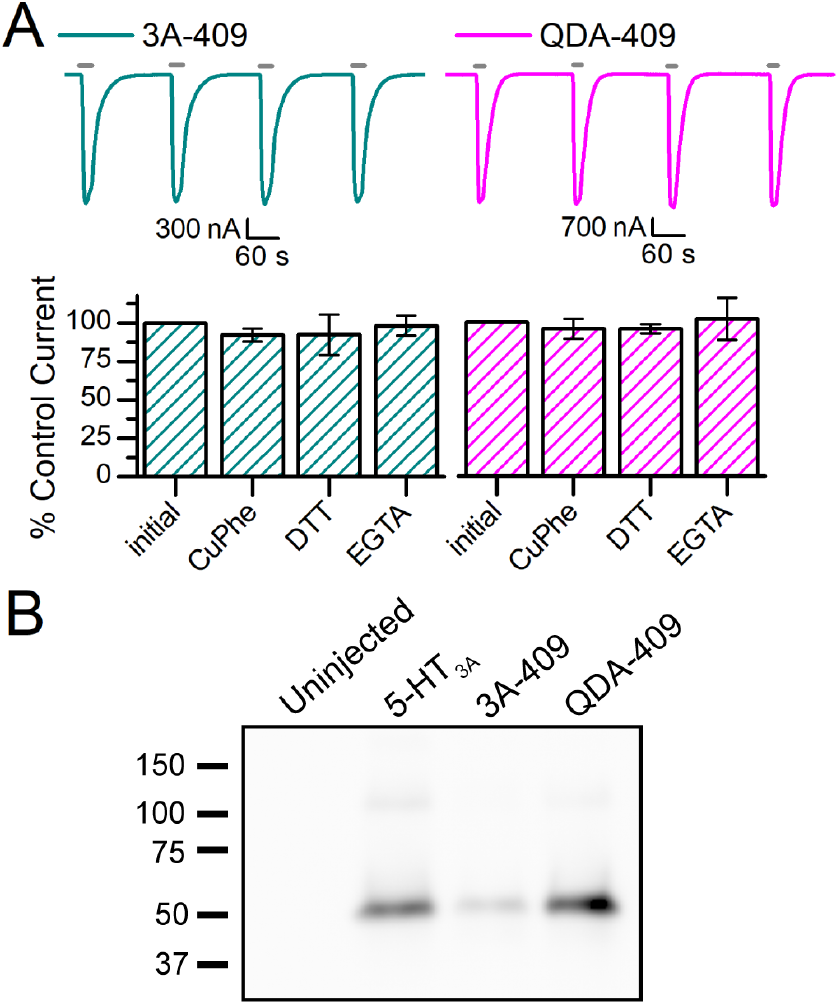
Evaluation of single Cys at I409 in response to 5-HT, Cu:Phe, DTT, and EGTA. **(A)** (top panel) Sample traces of 5-HT_3A_-I409C and QDA-I409C in response to 3 μM 5-HT, showing stable current amplitudes with each agonist application. (bottom panel) Quantification of current amplitudes after 5-HT_3A_-I409C and QDA-I409C were exposed to Cu:Phe (100:200 μM), 10 mM DTT, and 1mM EGTA. The first inward current recorded in response to 3 μM 5-HT is the reference current amplitude (100% of current). Experimental set-up was the same as in Fig. 3A. Data is shown as mean±S.D. (n≥3). Statistical significance was determined with one-way ANOVA, Dunnett’s multiple comparisons test between the initial inward current and each set of conditions. (B) Western blot of single Cys and uninjected oocytes and 5-HT_3A_ as controls. Experimental set-up was the same as in Fig. 2. Expressing *X. laevis* oocytes were exposed to 2 min of 100:200 μM Cu:Phe. SDS-PAGE fractions were run without reducing agent.

**Figure S3.**
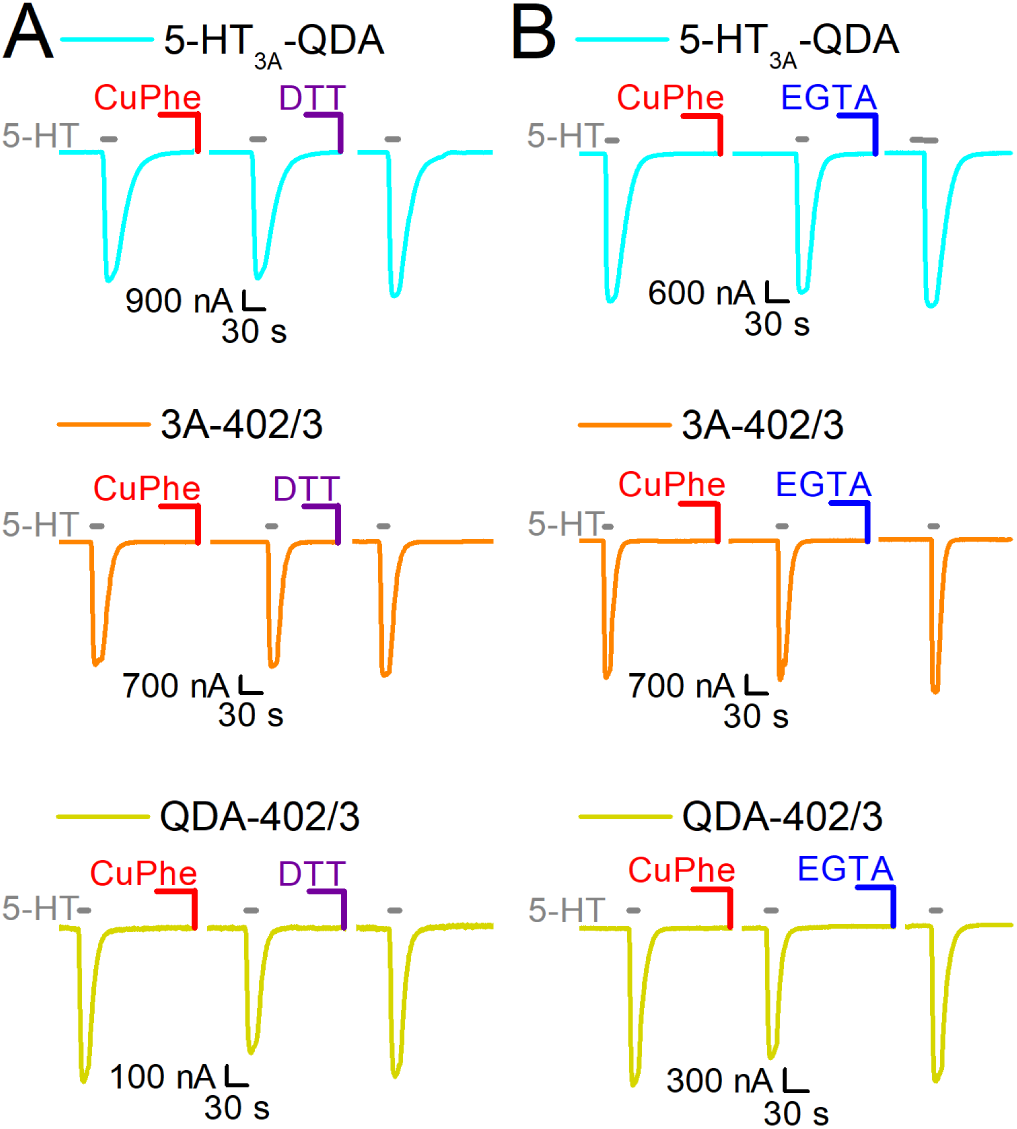
No Effect of Cu:Phe, DTT, and EGTA on QDA and L402C/L403C Cys pairs. **(A)** Sample traces of QDA (cyan), 5-HT_3A_-L402C/L403C (orange), and QDA-L402C/L403C (yellow). The inward currents were evoked with 3 μM 5-HT (grey bars). The initial inward current represents the reference current amplitude (100% of current). Following the 5-HT application(s), the oocytes were exposed to 2 min of Cu:Phe (100:200 μM) and then a 6 min wash with OR-2 (not pictured) before another application of 5-HT (second inward current depicted). 5-HT was applied until a stable current response was achieved (stable response to 5-HT is the inward current before DTT, labeled “5-HT” in Fig. 4 for I409C/R410C mutants). Lastly, 10 mM DTT was applied to the oocytes for 2 min, followed by a 6 min OR-2 wash (not pictured) and 5-HT (last inward current pictured). **(B)** Sample traces of QDA (cyan), 5-HT_3A_-L402C/L403C (orange), and QDA-L402C/L403C (yellow). Same experimental set-up as in **(A)** but during the last step 1 mM EGTA was applied instead of DTT.

